# FourCSeq: Analysis of 4C sequencing data

**DOI:** 10.1101/009548

**Authors:** Felix A. Klein, Simon Anders, Tibor Pakozdi, Yad Ghavi-Helm, Eileen E. M. Furlong, Wolfgang Huber

## Abstract

**Abstract:** *Motivation:* Circularized Chromosome Conformation Capture (4C) is a powerful technique for studying the spatial interactions of a specific genomic region called the “viewpoint” with the rest of the genome, both in a single condition or comparing different experimental conditions or cell types. Observed ligation frequencies show a strong, regular dependence on genomic distance from the viewpoint, on top of which specific interaction peaks are superimposed. Here, we address the computational task to find these specific interactions and to detect changes between interaction profiles of different conditions.

*Results:* We model the overall trend of decreasing interaction frequency with genomic distance by fitting a smooth monotonously decreasing function to suitably transformed count data. Based on the fit, *z*-scores are calculated from the residuals, with high *z* scores being interpreted as peaks providing evidence for specific interactions. To compare different conditions, we normalize fragment counts between samples, and call for differential contact frequencies using the statistical method `DESeq2` adapted from RNA-Seq analysis.

*Availability and Implementation:* A full end-to-end analysis pipeline is implemented in the R package `FourCSeq` available at www.bioconductor.org.

*Contact:*

## 2 Introduction

Circularized Chromosome Conformation Capture (4C) couples the low-throughput Chromosome Conformation Capture (3C) technique (Dekker *et al.*, 2002) for studying chromatin–chromatin interactions with high throughput sequencing (Simonis *et al.*, 2006; Stadhouders *et al.*, 2013). 4C detects the contacts of a chosen viewpoint with, in principle, the entire genome. The 4C protocol consists of six main steps (Stadhouders *et al.*, 2013). First, the chromatin is cross-linked with formaldehyde to fix DNA-protein complexes, thereby capturing DNA sequences that are in close spatial proximity. In the next step, the cross-linked chromatin is digested with a restriction enzyme. In the third step, the fragment ends from the digestion treatment are ligated under dilute conditions to favor intra-complex ligation, ligating DNA sequences that have been in close spatial proximity. After this, the cross-linking is reversed, followed by a second round of digestion with a different restriction enzyme to obtain smaller DNA molecules. These molecules are then circularized and PCR amplified. The resulting library is sequenced. The possibility to multiplex several viewpoints in one sequencing library further increases the throughput.

As result, the distribution of reads from a 4C sequencing library throughout the genome provides an estimate of the contact frequencies of the viewpoint with the rest of the genome. Overall, the 4C signal decreases with genomic distance from the viewpoint and reaches a constant level of noise for large distances. Specific interactions of DNA elements sit on top of this overall trend. The task is to extract interactions that stand out from the general trend. If 4C has been performed on samples with different cell types, developmental stages or experimental treatments, a possible next steps is the detection of changes in interaction frequencies between the sample groups.

Several analysis approaches for the first step, detection of interactions, have already been developed for 4C sequencing data. The approach by Thongjuea *et al.* (2013) uses a non-parametric smoothing spline on library-size normalized count data to estimate the signal decrease with distance to the viewpoint and detects interactions by calculating *z*-scores from the residuals of this fit. Another approach, used by van de Werken *et al.* (2012) and Splinter *et al.* (2012), is based on either a semi-quantitative contact profile in the proximity of the viewpoint, an empirically estimated contact background model or binary contact profile combined with a window based enrichment and permutation analysis.

Clearly missing is a method that uses replicate information to detect consistent peaks and to statistically infer changes in contact frequencies between different conditions.

We address these needs with the following approach. We use a distance-dependent monotonous fit to estimate the signal decay with increasing distance from the viewpoint, since the unspecific component of the signal decreases monotonously. As input to the fit we use variance-stabilized read count data (Anders and Huber, 2010). To detect strong interactions we calculate *z*-scores from the fit residuals and associated p-values. For the comparison of different conditions we use the methods implemented in the `DESeq2` package (Love *et al.*,*2014*

## 3 Materials and Methods

### 3.1 Data preprocessing

The data processing pipeline (Figure 1) starts from the reads of the 4C library. If several 4C libraries were multiplexed, the viewpoint primer sequences and, if present, additional barcodes, are used to demultiplex the sample and trim of the primer sequences. For the demultiplexing and trimming of primer sequences a Python script is included in the package. The remaining sequences are aligned to the full reference genome using a standard alignment tool.

**Figure 1:**
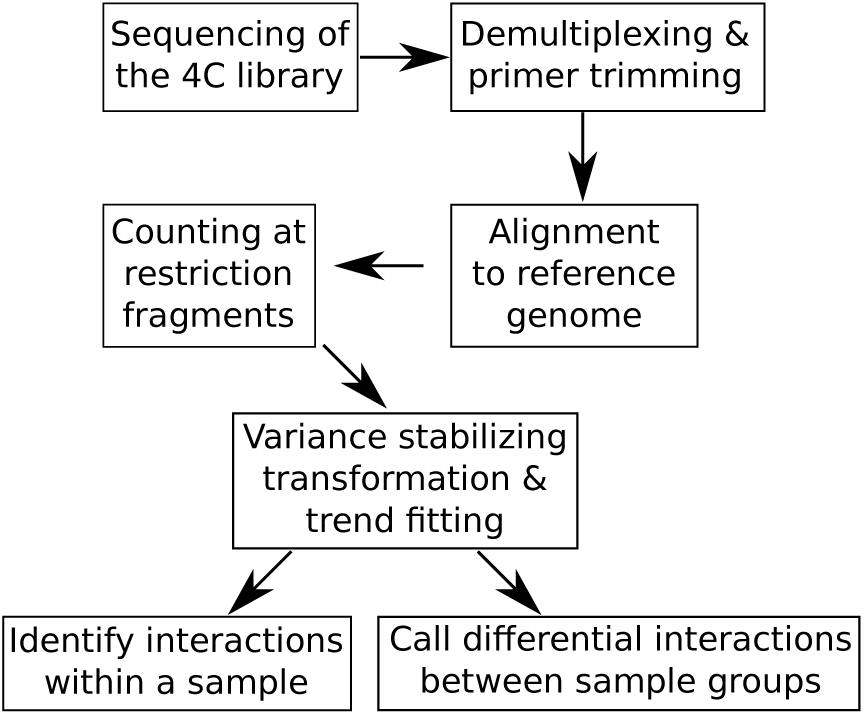
Overall workflow of steps described in this paper.

The analysis pipeline of our R package starts with the binary alignment/map (BAM) files output from the alignment. The following steps are now described in more detail.

#### 3.1.1 Cutting the reference genome

The input to the statistical analysis is a count table, with one row for each restriction fragment, and one column for each sample, with the table entries indicating how many reads have been assigned to each restriction fragment in each sample. By restriction fragment, we mean the sequences between the cutting sites of the first restriction enzyme, because this first digestion defines the resolution at which interactions can be seen in 4C. To define fragments, we cut the reference genome *in-silico* using the recognition sequence of the first cutter. Fragments are delimited by adjacent cutting sites of the first restriction enzyme. The second restriction enzyme is used to reduce the size of the fragments for efficient circularization and PCR amplification. Correspondingly, fragment ends are defined as the genomic region between the start/end position of the fragment and the cutting site of the second enzyme that is closest to the start/end position of the fragment respectively (Figure 2a).

**Figure 2:**
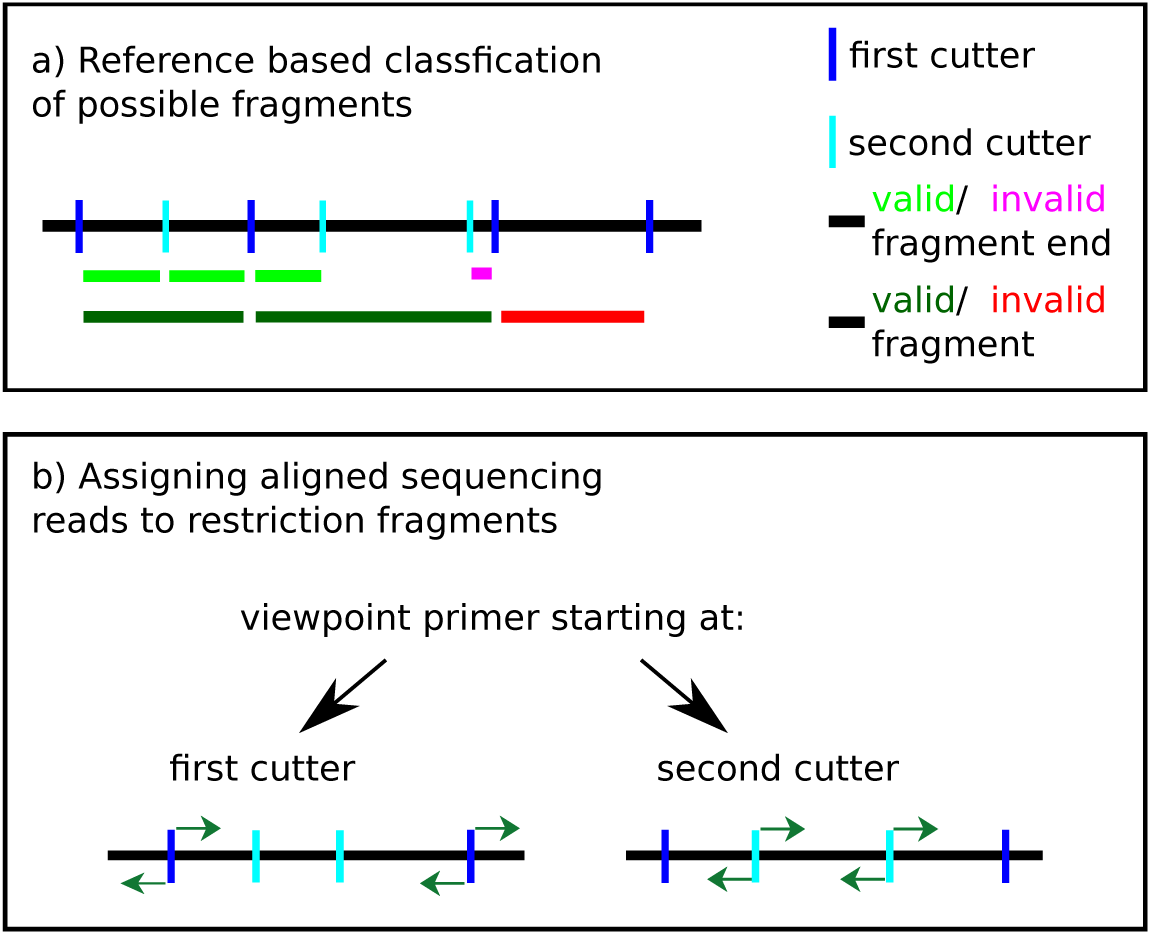
(a) Schematic of the rules to define valid fragments, i. e., fragments that are used subsequently in the analysis. The pink fragment end is smaller than the defined threshold, but since the other fragment end is valid, the fragment is kept for analysis. The red fragment is invalid because it does not contain a cutting site of the second restriction enzyme and is removed from the analysis. (b) If the sequencing primer starts at the first restriction enzyme cutting site, reads (green arrows) that start at the fragment ends and are oriented towards the fragment middle are kept for analysis. If the sequencing primer starts at the second restriction enzyme cutting site, reads (green arrows) that start right next to the cutting site of the second restriction enzyme and are directed towards the ends of the fragment are kept for analysis.

Since mainly fragments that contain a second cutting site are efficiently amplified by the protocol, a fragment is considered valid if it contains at least one cutting site of the second enzyme and has long enough fragments ends. By default, a threshold of 20 nt on the minimum length of a fragment end is used.

#### 3.1.2 Mapping of primer sequences

The primer sequence of the viewpoint is mapped to the reference genome to find the fragment that contains the viewpoint. This fragment is used to calculate the genomic distance to fragments on the same chromosome. The distance between the viewpoint and a fragment is taken as the genomic distance between the middle of the viewpoint fragment and the middle of the other fragment.

#### 3.1.3 Mapping reads to fragment ends

To filter out non-informative reads, we use the following criteria, motivated by the 4C protocol: Only reads that fulfill the criteria are mapped to a fragment end. A first condition is that reads should start directly at a restriction enzyme cutting site. Additionally, the orientation of the read at the fragment end is important and defined by the protocol (Figure 2b). If the sequencing library was prepared with a primer starting next to a cutting site of the first cutting enzyme, reads should be directed to the middle of the fragment. If the primer starts next to a cutting site of the second cutter instead, reads should be directed towards the fragment ends. The reads mapped to both fragment ends are combined for subsequent analysis. To check for consistency between replicates, scatter plots of count values can be created (Figure 3).

**Figure 3:**
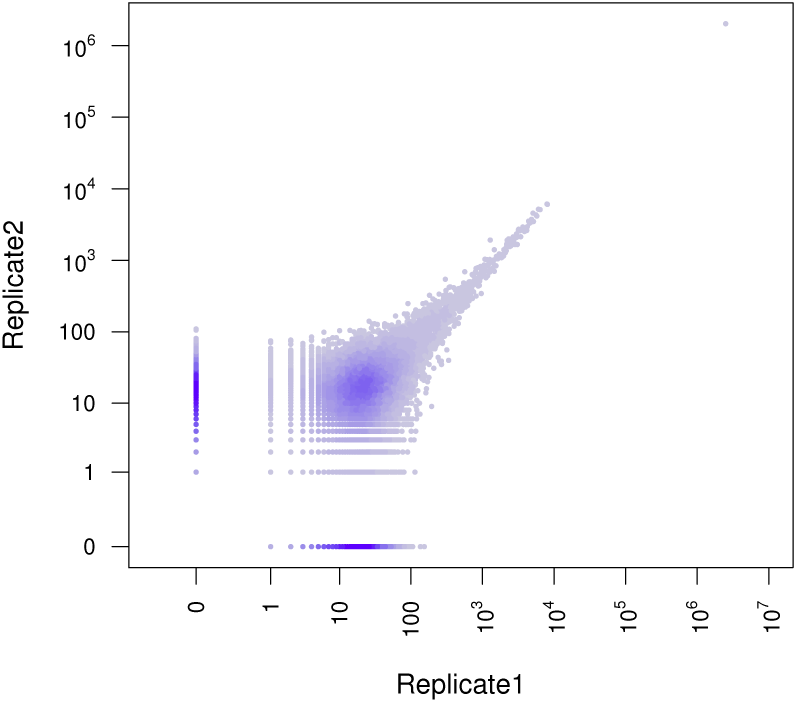
Correlation between two biological replicates of the *apterous* CRM viewpoint for whole embryo tissue at 6-8 h after fertilization. In the plot a desity estimate of the pairwise distribution of count values per fragment is shown. The *x*- and *y*-axes (drawn in logarithmic scale, with zero) correspond to the counts for the fragments in two biological replicate libraries for the same viewpoint and biological condition. The replicates show good concordance for higher count values. Fragments with 0 counts for both replicates are removed.

### 3.2 Detecting interactions

#### 3.2.1 Variance-stabilizing transformation

The count values usually span several orders of magnitude. If a logarithmic transformation were used for the count values, low abundance fragments would tend to show large standard deviations across samples. On the other hand, if untransformed data were used, the standard deviations across samples would be very large for high abundance fragments. This heteroscedasticity would skew the analysis towards either the fragments very far or very close to the viewpoint. Therefore, we use the variance-stabilizing transformation *v* as introduced by Anders and Huber (2010) and implemented in the `DESeq2` package (Love *et al.*, 2014), to transform the count *k*_*ij*_ of fragment *i* in sample *j* to *v*(*k*_*ij*_). After transformation the standard deviations show less dependence on the fragment abundance (Figure 4)

**Figure 4:**
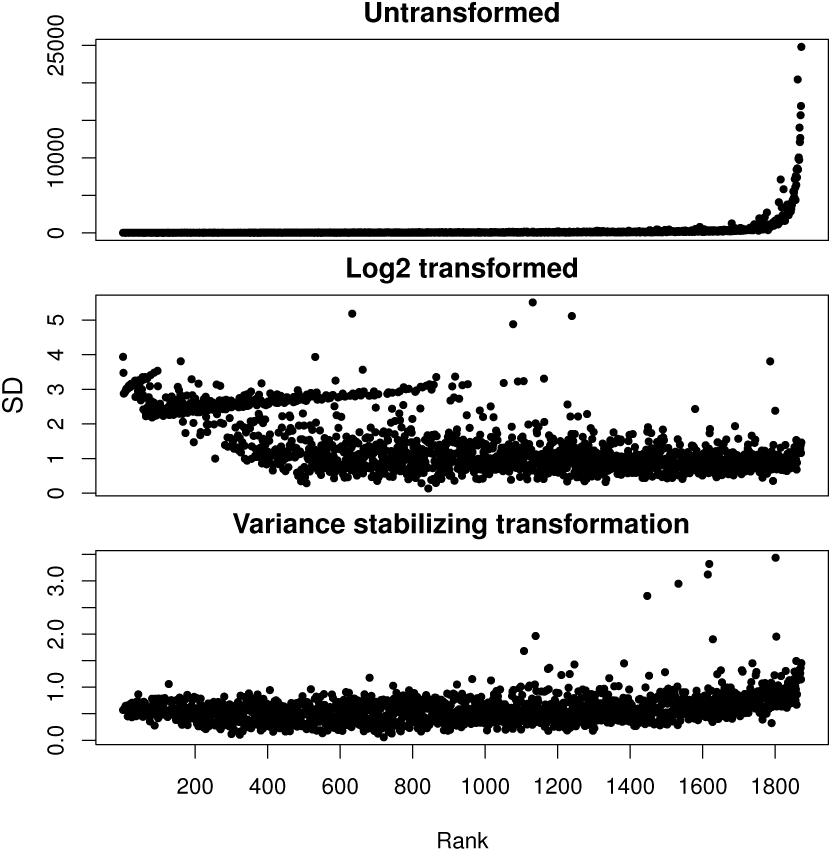
Variance-stabilizing transformation. For each fragment, the mean and standard deviation of its count data were computed across all samples for the *apterous* CRM viewpoint. The plots visualize the distributions of these values for all fragments. Fragments close to the viewpoint are on the right side with higher count values. When the untransformed count data are considered (upper panel), the standard deviations are very large for high abundance fragments (close to the viewpoint). When the count data are considered on the logarithmic scale (middle panel), the standard deviations are very large for low abundance fragments (far from the viewpoint). Both effects would make the analysis highly susceptible to noise either close or far from the viewpoint respectively. When the data are transformed using a variance stabilizing transformation, the standard deviations show less dependence on the fragment abundance, allowing for a more consistent statistical treatment across the whole dynamic range of the data.

#### 3.2.2 Trend fitting

The 4C signal decays with genomic distance from the viewpoint and converges towards a constant level of background. This decay trend *f*_*j*_(*d*_*i*_) is fitted using the transformed count values *v*(*k*_*ij*_) as a function of the logarithm of the genomic distance *d*_*i*_ from each fragment *i* to the viewpoint.

The FourC package offers two implementations for the distance dependence fit using the smooth monotone fit function of the fda package (Ramsay *et al.*, 2014). First, assuming that the general trend is symmetric around the viewpoint, we fit a symmetric monotone curve on the combined data from both sides. Second, we perform a monotone fit for each side of the viewpoint.

The latter case can be used if one is interested in finding asymmetries in the interaction profiles of a viewpoint, which might be of particular interest at boundaries of topological domains (Dixon *et al.*, 2012). For both methods we provide standard parameters that work for a wide range of data and which can be adjusted by the user if necessary.

An example of a symmetric monotone fit is shown in Figure 5.

**Figure 5:**
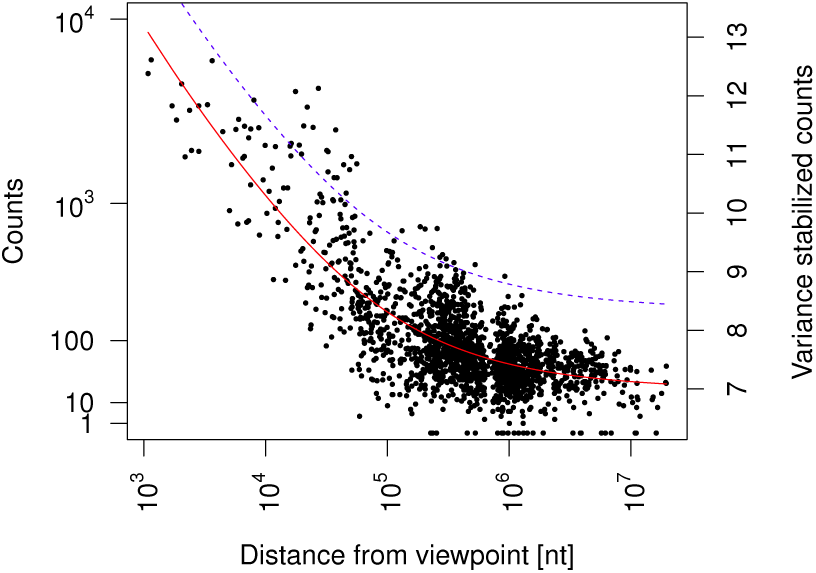
Symmetric monotonous fit of the variance-stabilized count data over the logarithm of the distance from viewpoint for the *apterous* CRM viewpoint. The red line shows the fit and the blue dashed line is the fit plus 3*σ*.

#### 3.2.3 *z*-scores of residuals

To find specific interactions that show higher interaction frequencies than expected at a given distance from the viewpoint, we calculate *z*-scores from the residuals of the fit and look for large positive *z*-scores. The *z*-scores are calculated in the following way:

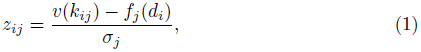

where *σ*_*j*_ = mad_*i*_(*v*(*k*_*ij*_) − *f*_*j*_(*d*_*i*_)) is a robust estimator of scale, *i* runs over all fragments and *j* over all samples.

Using the calculated *z*-scores and assuming that they follow a Normal distribution under the null hypothesis, one sided p-values are calculated for each fragment. These p-values are adjusted for multiple testing using the method of Benjamini and Hochberg (1995).

Specific interactions can now be found by looking for fragments with large positive *z*-scores and small adjusted p-values (Section 4.2).

### 3.3 Differences between conditions

We have observed the distance dependence of the signal to be variable between samples and this needs to be taken into account for comparisons. Therefore, we calculate a matrix of normalization factors *n*_*ij*_, such that the scaled read counts *n*_*ij*_ *k*_*ij*_ for fragment *i* become comparable across the samples *j*. To this end, we need the normalization factors to represent the fitted distance dependence on the scale of the raw counts. Hence, we back-transform the fitted values *f*_*ij*_ to the scale of raw counts and scale them to have unit geometric mean across samples to obtain the normalization factors:

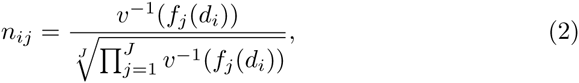

where *n*_*ij*_ is the normalization factor, *v*^-1^ (*f*_*j*_(*d*_*i*_)) is the back transformed fitted value at the genomic distance *d*_*i*_. The index *i* runs over all fragments and *j* over all samples.

With these normalization factors we use the methods implemented in the 

~~~
DESeq2
~~~

 package to detect differences between conditions (Love *et al.*, 2014). This approach performs a quantitative comparison of the normalized fragment counts for each single fragment between conditions. The fold-change between conditions is compared to the variability between biological replicates using a Wald test. Significant interaction changes are called when the observed change between conditions is significantly stronger than what is expected from the size of the changes seen between replicates.

## 4 Results

To illustrate our approach, we use a 4C data set of developing *Drosophila melanogaster* embryos (Ghavi-Helm *et al.*, 2014). In this data set 103 viewpoints were selected throughout the *D. melanogaster* genome focusing on *cis*-regulatory modules (CRMs). Samples were taken from embryos at 2-4 h and 6-8 h after fertilization either using whole embryos or mesoderm-specific cells (Ghavi-Helm *et al.*, 2014).

### 4.1 Preprocessing

Starting from FASTQ, files we used a Python program, which is included in the `FourCSeq` package, to demultiplex the libraries and trim off bar codes and adapters. Next we aligned the reads to the dm3 reference genome with Novoalign (http://www.novocraft.com).

For short restriction fragments, we observed the problem that reads contained the whole fragment and then continued through the cutting site of the second restriction enzyme into the ligated fragments (in most cases the viewpoint fragment). This often resulted in two possible alignments causing the reads to be reported as not uniquely mapping. To address this problem and rescue some of the shorter fragments we checked whether the restriction enzyme cutting site was found within unaligned reads. In such a case the end of the read was trimmed at the restriction enzyme cutting site and alignment was attempted again.

We then generated a fragment reference and mapped the aligned reads to these fragments as described in Sections 3.1.1 and 3.1.3.

For quality control, the percentage of reads mapping to valid fragments from all aligned reads was calculated. For our data this value was around 70-95% in most cases. A value in that range should be obtained for a 4C library. If the percentage is much smaller, the first region that should be investigated is the region around the viewpoint, where a single fragment can pile up a high percentage of the reads. Other possible reasons for low mapping percentages might be reads that map either to invalid fragments, which have been removed from analysis, or to new fragments, created by mutations relative to the reference genome.

To check whether technical and biological replicates gave a similar signal, scatter plots of the replicates were generated. For our data set, these plots showed good agreement for higher count values in most cases. However, at lower count values, the replicates show higher relative variation, as is expected from Poisson noise. An example is shown in Figure 3.

### 4.2 Detecting interactions

First, to reduce noise in the data, we removed fragments which had less than a median of 40 counts across all samples for one viewpoint. Second, we removed fragments that were too close to the viewpoint, because the high signal obtained from these fragments is generally due to ligation caused by close linear proximity to the viewpoint. The package therefore automatically defined the first valid fragments as those that occurred directly after the strong initial signal decrease with distance from the viewpoint where the signal began to increase again. The parameters of the variance-stabilizing transformation were fitted on the count values of the remaining fragments.

The result of the variance-stabilizing transformation is shown in Figure 4. Next the decay trend is fitted on the transformed scale, using a monotonous symmetric fit. The fit is shown in Figure 5. *z*-scores and associated p-values were calculated from the fit residuals. Interactions were found by looking for fragments with *z*-scores larger than 3 in both replicates and an adjusted p-values smaller than 0.01 in at least one replicate. Figure 7 shows the results for one of the viewpoints in ou data set, which is located in a cis-regulatory module (CRM) close to the *apterous* (*ap*) gene. The fragments that show an interaction are highlighted by red dots.

In mesoderm specific and whole embryo tissue at 6-8 h after fertilization the interaction of the viewpoint with the *ap* gene promoter on the right side of the viewpoint is captured. Further interactions are found as well, but could not be directly attributed to a specific genomic element. In general we were able to detect interactions between ten known enhancer-promoter pairs and many more interactions throughout the set of 103 viewpoint (Ghavi-Helm *et al.*, 2014).

### 4.3 Differences between conditions

To detect differences between conditions, we used the method described in Section 3.3.

Figure 6 shows the MA plot comparing mesoderm tissue and whole embryo for *Drosophila* embryos 6-8 h after fertilization. For the same viewpoint the results of the analysis are shown in Figure 7. Fragments that have a adjusted p-value of less than 0.01 for the Wald test are highlighted by blue points, or by orange points, if they additionally are called as an interaction in the depicted sample.

**Figure 6:**
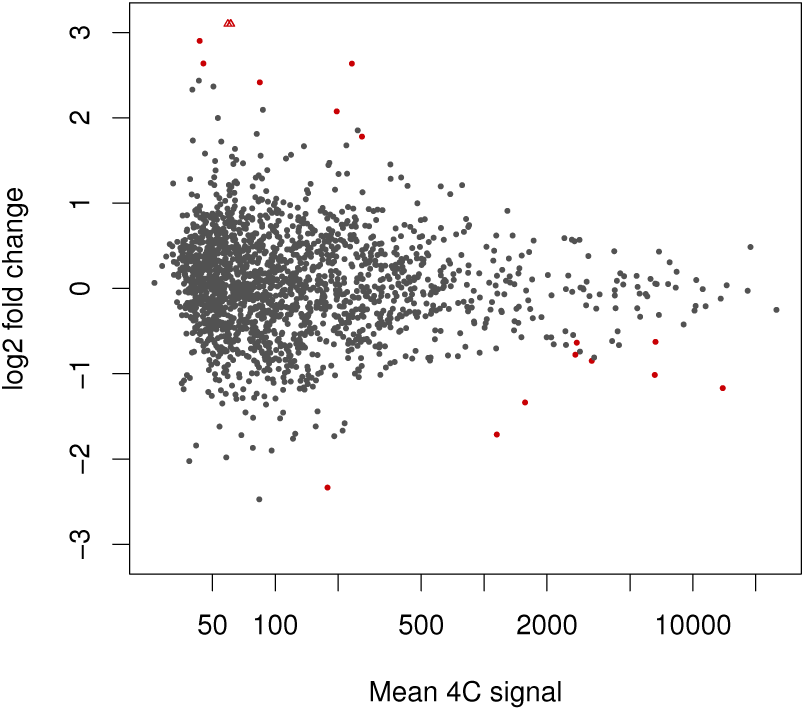
MA plot of the *apterous* CRM viewpoint 4C profile comparison between *Drosophila* embryo mesoderm tissue and whole embryo 6-8 h after fertilization. The y axis shows the difference between log interaction counts for a given fragment which is plotted against the average log interaction counts per fragment on the x-axis. Red dots represent fragments that show differential interactions (p-adjusted *<* 0.01)

**Figure 7:**
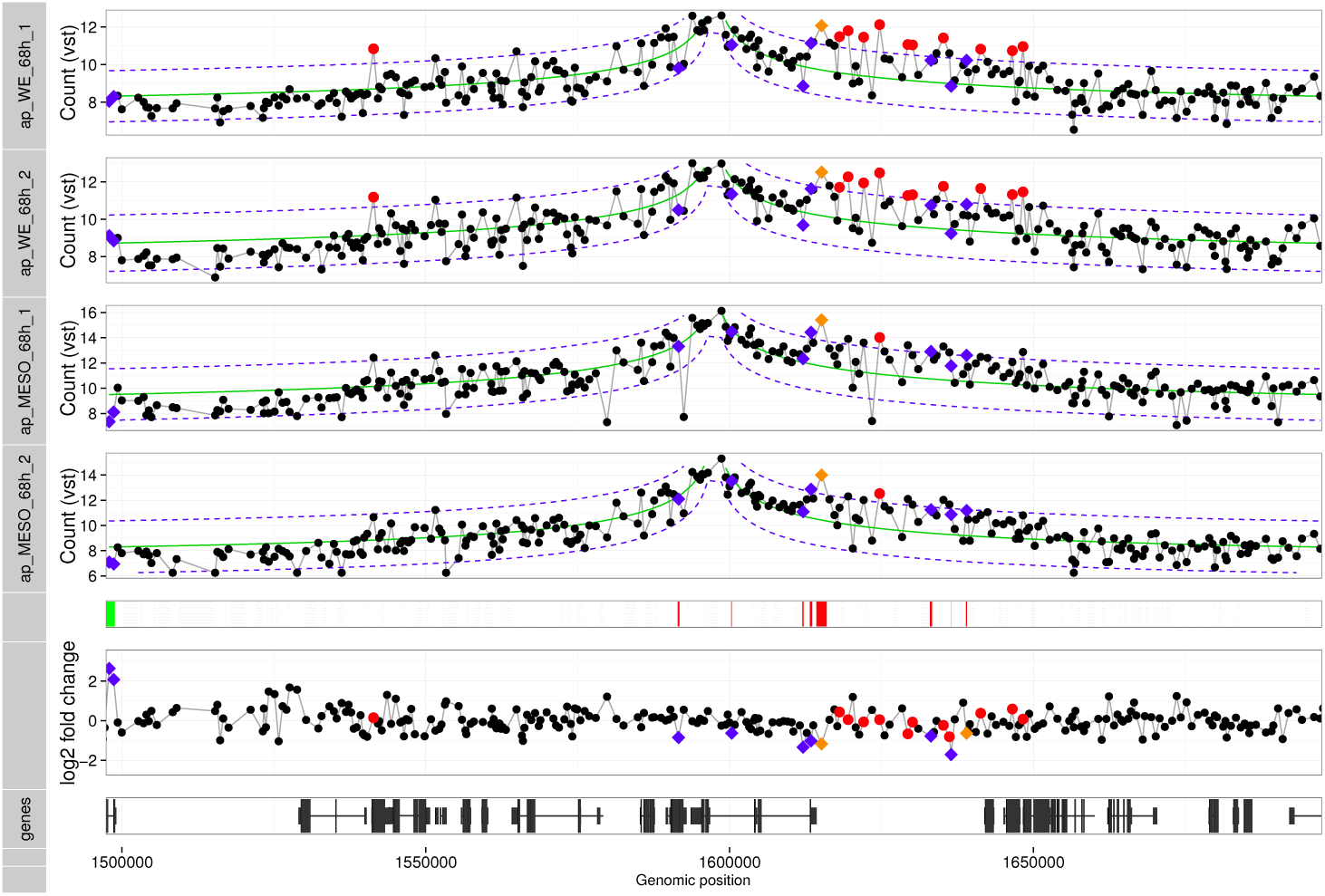
Detection of interactions and differences: The figure shows the plot generated by the `FourCSeq` to visualize the results. The upper 4 wide tracks show the variance-stabilized counts for 2 biological replicates of *Drosophila* embryo mesoderm tissue and whole embryo 6-8 h after fertilization for the *apterous* CRM viewpoint. The fit of the distance dependence is shown as solid green line and the dashed blue lines represent the fit ±3*σ*. Interactions detected by *z*-score *>* 3 in both replicates and p-adjusted *<* 0.01 for at least one replicate are shown as red or orange points. Fragments represented by orange points additionally show a differential interactions (p-adjusted *<* 0.01, differential Wald test). Differential changes in the contact profile that are not called as interactions are shown as blue points (p-adjusted *<* 0.01, differential Wald test). The color bar below the 4C profiles shows whether the upper condition (green) or the lower condition (red) has the higher signal for the detected differences (p-adjusted *<* 0.01). The calculated log2 fold-change of the differential testing per fragment are shown above the track at the bottom, which shows the gene model of the region.

In general one can observe that the effect sizes for differential changes are very small and the overall pattern of the interaction profiles remains largely unchanged, as we recently reported (Ghavi-Helm *et al.*, 2014).

However, for the strong interaction at the *ap* promoter we estimate a significant fold change of 2.25 between the conditions. Stronger contacts in the mesoderm tissue could be due to the fact that the ap gene is only expressed in the mesoderm. Only 6 % of identified interactions showed evidence of interaction changes across time and tissue context (Ghavi-Helm *et al.*, 2014).

## 5 Discussion

We described the functionality of the `FourCSeq` package available on Bioconductor.

Our approach to detect peaks is in general similar to the method implemented in the `r3cseq` package (Thongjuea *et al.*, 2013). However, while Thongjuea *et al.* (2013) do the fit on raw count scale, we use a variance-stabilizing transformation on the data to have a more consistent statistical treatment across the different orders of magnitude observed in the count data. To detect specific interactions, we fit the decay of the variance stabilized 4C signal with distance from the viewpoint and calculate *z*-scores from the fit residuals. With this more consistent approach we were able to detect long-range chromatin interactions that span genomic distances *>* 100 kb in the compact *Drosophila* genome (Ghavi-Helm *et al.*, 2014).

Instead of only looking at log2 fold-changes of single samples between conditions as it is done in `r3cseq`, we make use of the framework for differential expression analysis implemented in the `DESeq2` package to detect differences between groups of samples in different experimental conditions. With this approach we take the variability between replicates of the data for each genomic position into account for the quantitative comparison of the fragment counts between conditions. The fold-change between conditions is compared to the variability of the data between biological replicates, and differential interaction are called statistically significant only if the observed fold change between conditions is significantly higher than what it is expected based on the noise level in the data.

In contrast to our method, the method by van de Werken *et al.* (2012) uses a customized approach for aligning reads to a reference of fragment ends. This alignment data can be further normalized and visualized by the tool that they provide. The results are plots of contact profiles and contact domainograms generated by analyzing the data with different window sizes. However, with this approach, comparisons of interaction profiles are only made qualitatively.

Our implementation allows the use of any FASTA file as reference genome; for example the dm3 genome was used for the data shown in Section 4, whereas the `r3cseq` package is limited to the mm9, hg18 and hg19 genomes.

To integrate called interactions and differences with other genomic data the results from our package can be used within the Bioconductor framework of *GenomicRanges* (Lawrence *et al.*, 2013). Furthermore we provide the possibility to export the interaction profiles as bigWig files for visual inspection in a genome browser along with other tracks of interest.

In summary, our package provides the tools to analyze 4C sequencing data and integrate the results with other genomic features. Its use will help to further investigate and understand the role of chromatin 3D structure in biological processes such as gene regulation and embryogenesis.

## Acknowledgments

The authors thank the members of the Huber group for discussion and comments, in particular A. Pekowska and B. Klaus for their valuable suggestions. *Funding:* F.A.K., S.A. and W.H were supported by the EC FP7 project Radiant. TP was supported by an ITN grant (EvoNet) and DFG grant (FU750).E.E.M.F. was supported by a DFG (FU 750) grant and Y.G.-H. by an EMBO post-doctoral fellowship.

## Conflict of Interest

none declared.

